# Opposing effects of T cell receptor signal strength on CD4 T cells responding to acute versus chronic viral infection

**DOI:** 10.1101/2020.08.06.236497

**Authors:** Marco Künzli, Peter Reuther, Daniel D. Pinschewer, Carolyn G. King

**Author notes:** To whom correspondence should be addressed: Carolyn King, Department of Biomedicine, University of Basel, University Hospital Basel, Hebelstrasse 20, CH-4031 Basel, Switzerland.

## Abstract

A hallmark of the adaptive immune response is the ability of CD4 T cells to differentiate into a variety of pathogen appropriate and specialized effector subsets. A long-standing question in CD4 T cell biology is whether the strength of TCR signals can instruct one Th cell fate over another. The contribution of TCR signal strength to the development of Th1 and T follicular helper (Tfh) cells has been particularly difficult to resolve, with conflicting results reported in a variety of models. Although cumulative TCR signal strength can be modulated by the infection specific environment, whether or not TCR signal strength plays a dominant role in Th1 versus Tfh cell fate decisions across distinct infectious contexts is not known. Here we characterized the differentiation of CD4 TCR transgenic T cells responding to a panel of recombinant wild type or altered peptide ligand lymphocytic choriomeningitis viruses (LCMV) derived from acute and chronic parental strains. We found that while TCR signal strength positively regulates T cell expansion in both infection settings, it exerts opposite and hierarchical effects on the balance of Th1 and Tfh cells generated in response to acute versus persistent infection. The observation that weakly activated T cells, which comprise up to fifty percent of an endogenous CD4 T cell response, support the development of Th1 effectors highlights the possibility that they may resist functional inactivation during chronic infection. We anticipate that the panel of variant ligands and recombinant viruses described herein will be a valuable tool for immunologists investigating a wide range of CD4 T cell responses.

**Graphical abstract:** 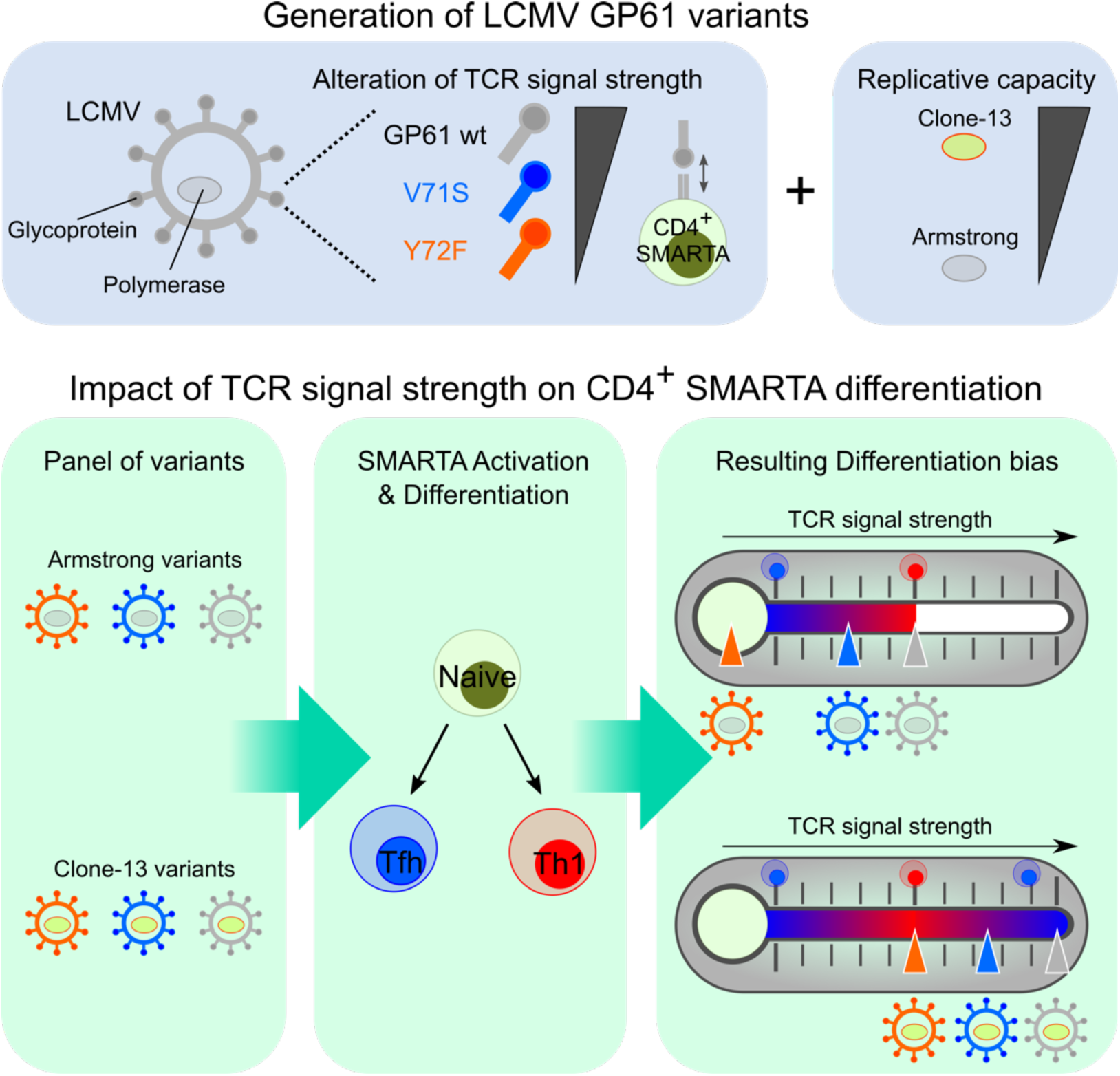

**Highlights:**

- Identification of a wide panel of altered peptide ligands for the LCMV-derived GP61 peptide
- Generation of LCMV variant strains to examine the impact of TCR signal strength on CD4 T cells responding during acute and chronic viral infection
- The relationship between TCR signal strength and Th1 differentiation shifts according to the infection context: TCR signal strength correlates positively with Th1 generation during acute infection but negatively during chronic infection.

## Introduction

Following infection or vaccination, antigen specific T cells undergo clonal expansion and differentiation into effector cells with specialized functions. This process begins with T cell receptor (TCR) recognition of peptide/MHC (pMHC) on antigen presenting cells (APCs) and is further modulated by cytokines and costimulatory molecules(*1, 2*). Viral infection induces the early bifurcation of CD4 T cells into Th1 and T follicular helper (Tfh) cells. Th1 cells potentiate CD8 T cell and macrophage cytotoxicity, whereas Tfh cells support antibody production by providing survival and proliferation signals to B cells(*1, 3*).

Although the cumulative strength of interaction between TCR and pMHC has a clear impact on T cell expansion and fitness, its influence on the acquisition of Th1 and Tfh cell fates is controversial(*4-12*). An essential role for TCR signal was implicated in a study assessing the phenotype of progeny derived from individual, TCR transgenic (tg) T cells responding during infection(*9*). The authors observed that distinct TCRs induced reproducible and biased patterns of Th1 and Tfh phenotypes. Although earlier reports suggested that Tfh cell differentiation requires high TCR signal strength, recent work supports the idea that Tfh cells develop across a wide range of signal strengths, while increasing TCR signal intensity favors Th1 generation(*4-11*). A central difficulty in reconciling these findings is the use of different TCR tg systems as well as immunization and infection models that may induce distinct levels of costimulatory and inflammatory signals known to influence T cell differentiation. Although existing reports suggest that persistent TCR signaling drives a shift towards Tfh differentiation during chronic infection, whether this outcome can be modulated by TCR signal strength has not been examined(*13-15*).

The impact of TCR signal strength on CD4 T cell differentiation *in vivo* has been historically challenging to address. The use of MHC-II tetramers to track endogenous polyclonal T cell responses does not adequately detect low affinity T cells that can comprise up to fifty percent of an effector response in autoimmune or viral infection settings(*16*). TCR tg models paired with a panel of ligands with varying TCR potency have been informative, but only a handful of MHC-II restricted systems exist(*17, 18*). To bypass these limitations, the generation of novel transgenic/retrogenic TCR strains or recombinant pathogen strains is required(*4, 12, 19, 20*). To our knowledge, this approach has not yet been used to modify a naturally occurring CD4 T cell epitope of an infectious agent. To test the impact of TCR signal strength across different types of infectious contexts, we generated a series of lymphocytic choriomeningitis virus (LCMV) variants by introducing single amino acid mutations into the GP61 envelope glycoprotein sequence and expressing them from both acute and chronic parent strains. These strains were used to assess the dynamics and differentiation of SMARTA T cells, a widely used TCR tg mouse line that mirrors the endogenous, immunodominant CD4 T cell response to LCMV(*13*). We observed that depending on the infection setting, TCR signal strength has opposing effects on the balance between Th1 and Tfh cell differentiation. In an acute infection, strong TCR signals preferentially induce Th1 effectors, whereas weak TCR signals shift the balance toward Tfh effectors. In contrast, strong T cell activation during chronic infection induces Tfh cell differentiation while more weakly activated T cells are biased to differentiate into Th1 cells. Based on these findings we propose a Goldilocks model for the generation of Th1 effectors during viral infection, where too little or too much TCR signaling skews the CD4 T cell response toward Tfh differentiation.

## Results

### Generation and viral fitness of GP61 LCMV variants

To generate recombinant LCMV variants, we first screened a panel of altered peptide ligands (APLs) with single amino acid mutations in the LCMV derived GP61 peptide. Using the early activation marker CD69 as a proxy for TCR signal strength we identified 75 APLs for the SMARTA TCR transgenic line (Figure S1A, Figure 1A, B, Table 1). We selected twelve of these APLs, covering a wide range of T cell activation potential, to generate recombinant variant viruses, using site-directed PCR mutagenesis (*21*). Five APL-encoding sequences were successfully introduced into the genomes of both LCMV Armstrong (Armstrong variants) and Clone-13 (Clone-13 variants), the latter of which contains a mutation in the polymerase gene L that enhances the replicative capacity of the virus, enabling viral persistence(*22, 23*). To exclude a potential impact of differential glycoprotein-mediated viral tropism on CD4 T cell differentiation, we equipped both the Armstrong- and Clone-13-based viruses with the identical glycoprotein of the WE strain and introduced the epitope mutations therein (resulting viruses referred to as rLCMV Armstrong and rLCMV Clone-13, respectively)(*24*). Of these viruses, two variants, V71S and V72F (EC50 ∼ 0.1 µM and 1µM, respectively) demonstrated comparable viral fitness *in vitro* and *in vivo* (Figure 1C, D). We further performed an out-competition assay with invariant chain knockout splenocytes to ensure comparable presentation of these APLs by MHC-II (Figure 1E)(*25*). Taken together, these data demonstrate the development of a novel tool to examine the impact of TCR signal strength on SMARTA T cells activated by either acute or chronic viral infection.

**Table 1:** APLs with altered potential to activate SMARTA and corresponding EC_50_ values.

| APL | EC50 [M] | APL | EC50 [M] |
| --- | --- | --- | --- |
| GP61wt | 5.2E-09 | V71 W | 5.9E-08 |
| Y68 A | 1.3E-06 | V71 P | 2.2E-06 |
| Y68 V | 7.2E-08 | V71 M | 1.2E-08 |
| Y68 L | 6.5E-06 | V71 C | 2.1E-07 |
| Y68 I | 1.3E-07 | Y72 S | 4.2E-05 |
| Y68 S | 2.6E-06 | Y72 T | 1.7E-05 |
| Y68 T | 4.8E-07 | Y72 E | 2.1E-07 |
| Y68 N | 1.6E-07 | Y72 R | 2.1E-06 |
| Y68 E | 7.0E-05 | Y72 F | 9.6E-07 |
| Y68 Q | 6.7E-06 | Y72 P | 6.3E-08 |
| Y68 K | 1.1E-06 | Y72 M | 5.6E-05 |
| Y68 R | 1.5E-06 | Q73 L | 8.7E-08 |
| Y68 H | 3.1E-07 | Q73 S | 2.5E-05 |
| Y68 W | 1.4E-08 | Q73 D | 3.2E-06 |
| Y68 P | 1.1E-05 | Q73 E | 3.1E-07 |
| Y68 C | 1.0E-06 | Q73 K | 8.3E-06 |
| Y68 M | 1.6E-07 | Q73 W | 2.6E-07 |
| K69 G | 5.2E-06 | Q73 F | 1.1E-07 |
| K69 A | 3.2E-08 | Q73 C | 2.9E-07 |
| K69 V | 5.0E-07 | F74 V | 8.7E-07 |
| K69 L | 5.5E-07 | F74 I | 1.9E-05 |
| K69 S | 1.2E-07 | F74 Y | 1.3E-06 |
| K69 T | 8.1E-08 | K75 R | 5.1E-07 |
| G70 V | 3.5E-07 | S76 D | 1.0E-06 |
| G70 L | 1.2E-07 | S76 E | 5.7E-07 |
| G70 I | 3.5E-06 | S76 Q | 5.9E-08 |
| G70 K | 5.8E-08 | S76 K | 2.0E-07 |
| G70 R | 1.1E-07 | S76 R | 2.2E-06 |
| G70 F | 1.4E-05 | S76 W | 7.9E-05 |
| G70 Y | 1.7E-07 | S76 F | 4.5E-07 |
| G70 M | 6.2E-08 | S76 Y | 5.3E-06 |
| V71 G | 9.3E-07 | S76 P | 3.8E-07 |
| V71 S | 1.4E-07 | S76 C | 5.9E-08 |
| V71 D | 1.4E-06 | V77 G | 1.8E-07 |
| V71 N | 8.4E-06 | V77 S | 3.6E-08 |
| V71 E | 9.5E-06 | V77 D | 4.8E-07 |
| V71 Q | 2.9E-06 | V77 E | 6.5E-07 |
| V71 H | 2.2E-08 | V77 P | 4.0E-08 |

**Figure 1:**
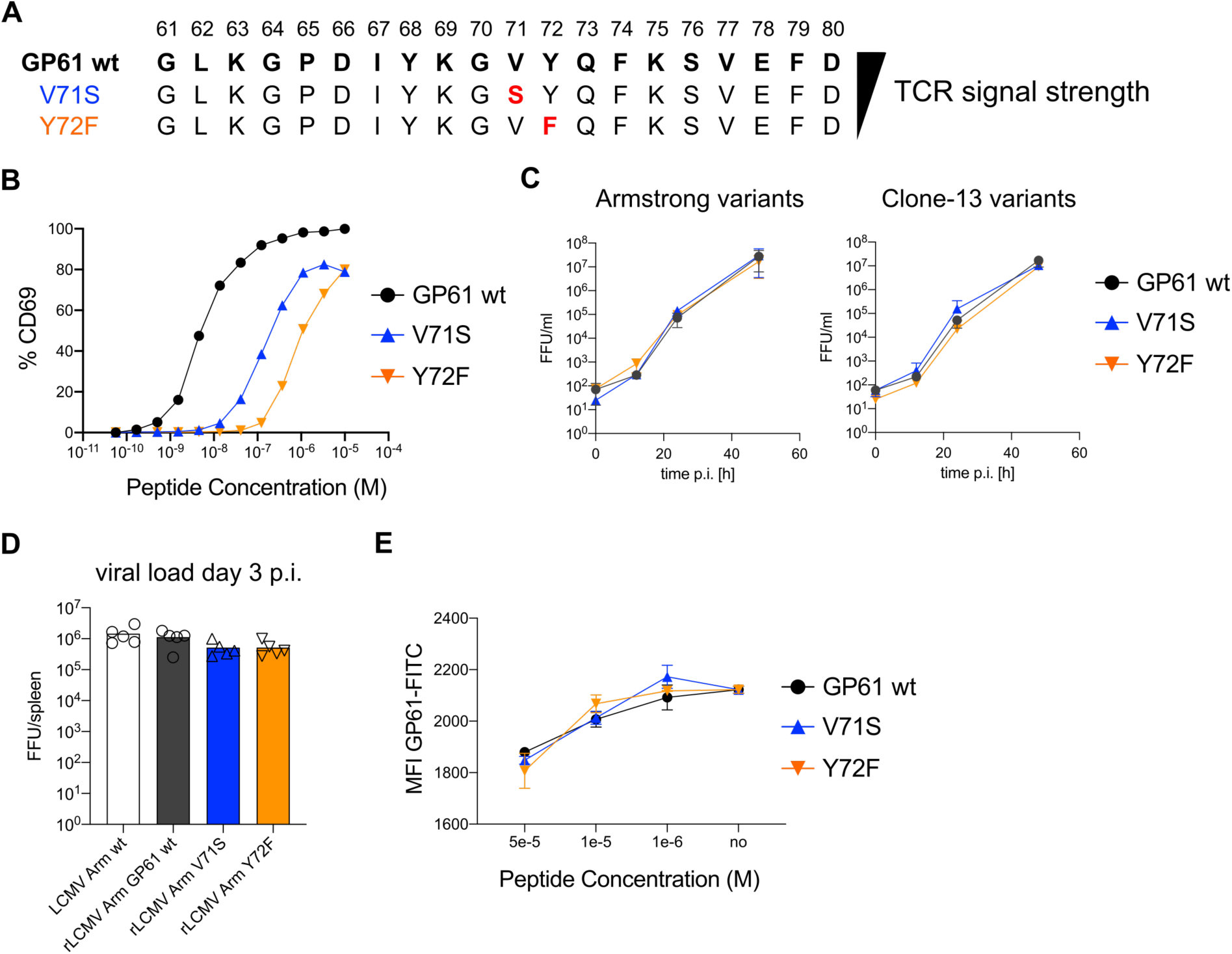
Generation and viral fitness of GP61 LCMV variants. (A) Scheme of GP61 wt and APL sequences with mutations highlighted in red ordered hierarchically according to TCR signal strength. (B) Peptide dose – activation curves of overnight cultured SMARTA cells with peptide pulsed splenocytes using the percentage of CD69^+^ SMARTA cells as a readout for activation. EC_50_ values are ∼ 5 nM for GP61 wt, ∼ 0.1 µM for V71S and ∼ 1µM for Y72F. (C) *In vitro* growth kinetics depicting the viral load in the culture medium (FFU/ml, focus forming units) of GP61 wt or V71S and Y72F variants of Armstrong (left) and Clone-13 (right) variant infection on BHK21 cells over time. Data are displayed as mean ± SD. (D) Early splenic viral load day 3 post infection (p.i.) in Armstrong variants. Bars represent the mean and symbols represent individual mice. (E) Peptide dose – response curves depicting the out-competition of the GP61 FITC signal by unlabeled GP61 wt or variants on B220^+^ B cells. Data are displayed as mean ± SD of 2-3 technical replicates. Data represent one of n = 2 independent experiments (B, D-,E) or pooled data from n = 2 independent experiments (C).

**Figure S1:**
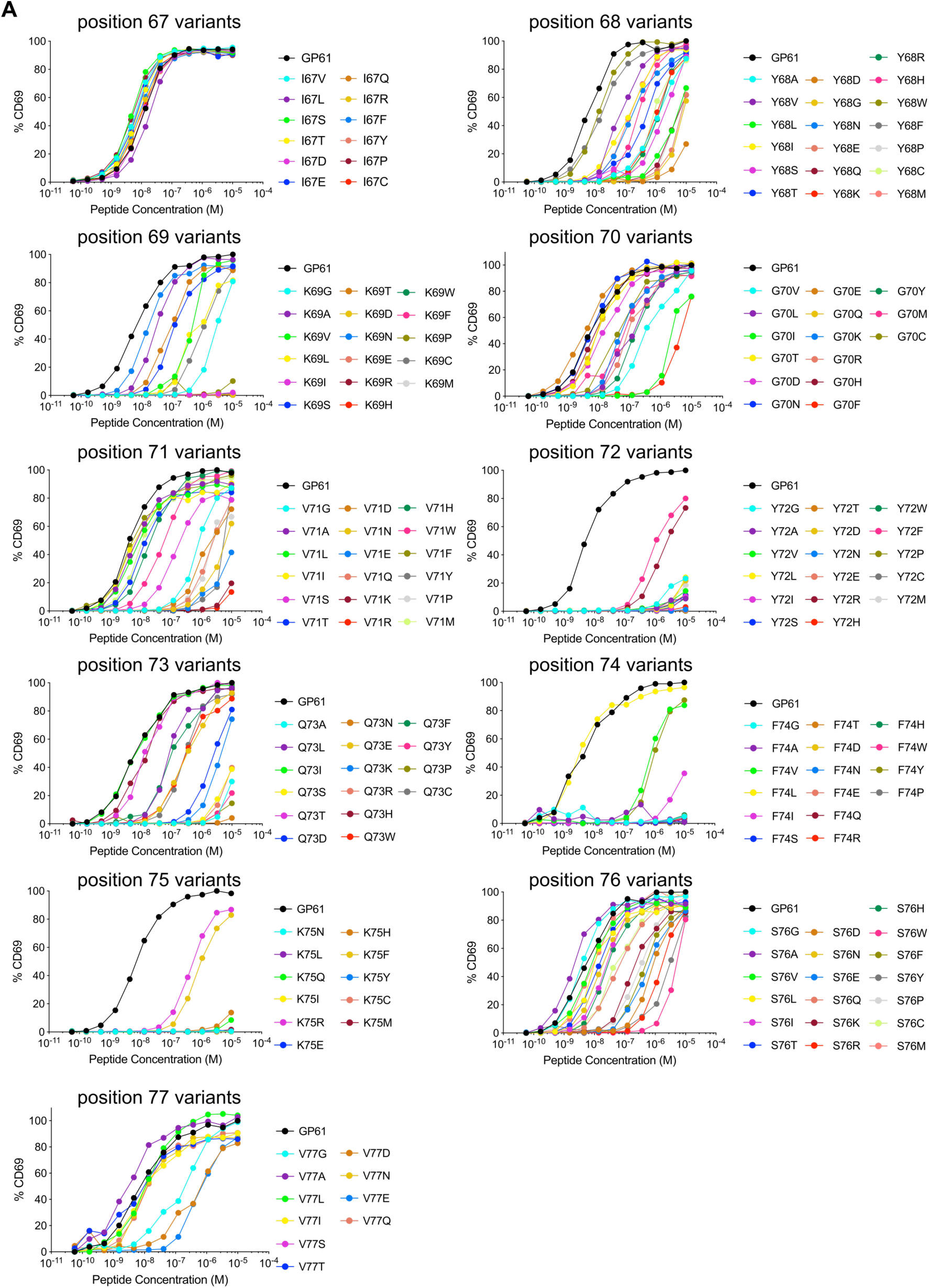
Generation and viral fitness of GP61 LCMV variants. (A) Peptide dose – activation curves of overnight cultured SMARTA cells with peptide pulsed splenocytes using the percentage of CD69^+^ SMARTA cells as a readout for activation. Data represent one of n = 2 independent experiments.

### TCR signal strength positively correlates with Th1 cell differentiation during LCMV Armstrong variant infection

To assess the impact of TCR signal strength during acute viral infection, SMARTA T cells were transferred into congenic recipients followed by infection with rLCMV Armstrong GP61 wt, V71S or Y72F. All LCMV variants were capable of inducing SMARTA T cell expansion at day 10 post infection (p.i.) and a direct correlation between TCR signal strength and the number of SMARTA T cells recovered was observed (Figure 2A). In contrast, expansion of endogenous LCMV nucleoprotein (NP)-specific as well as antigen-experienced CD44^+^ T cells was similar across all three viral strains (Figure 2A). The expansion hierarchy among the viruses was maintained >30 days after LCMV infection (Figure 2A).

**Figure 2:**
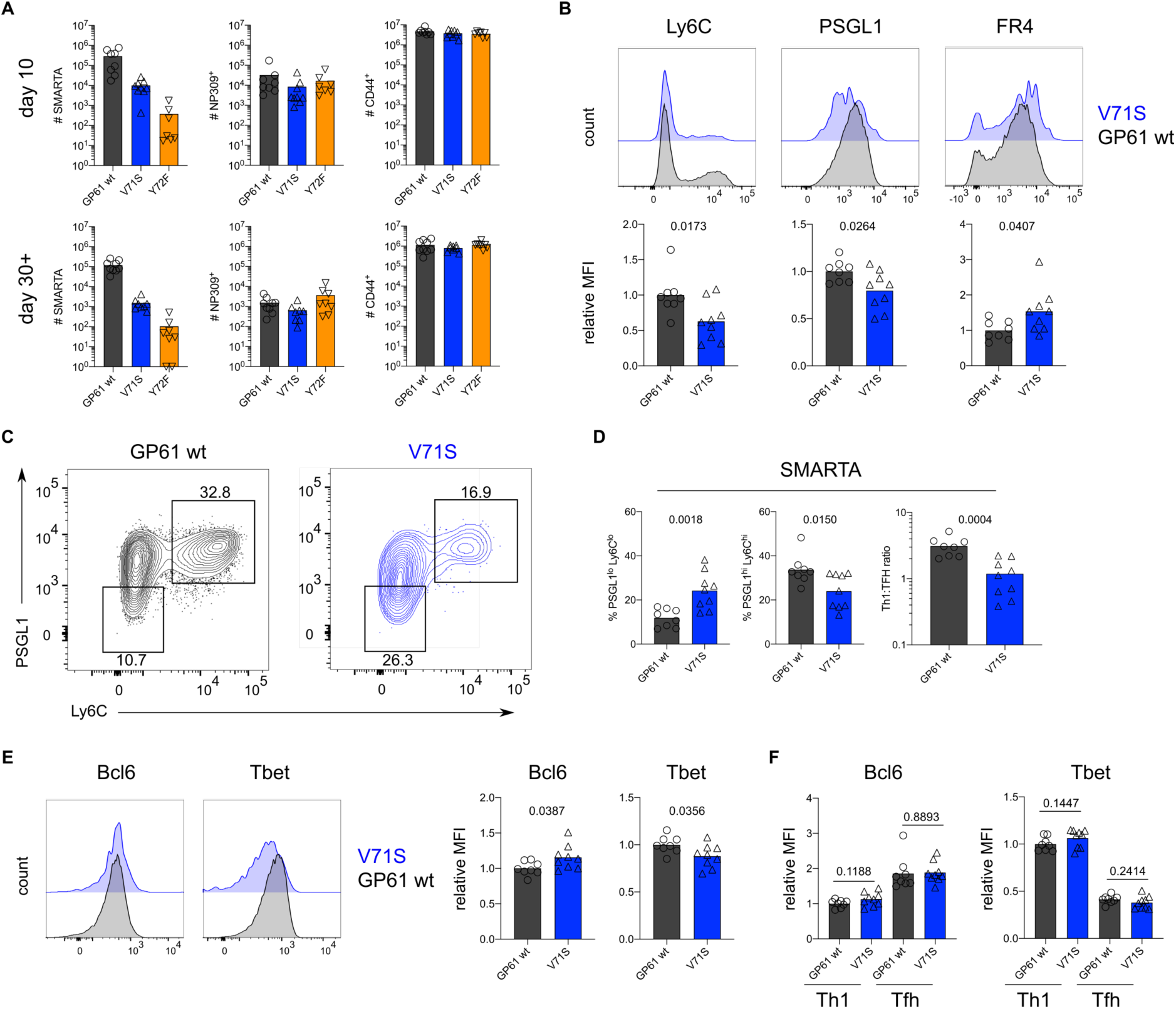
TCR signal strength positively correlates with Th1 cell differentiation during LCMV Armstrong variant infection. (A) Number of SMARTA (left), NP309^+^ (middle) and CD44^+^ cells (right) 10 days (top) or >30 days (bottom) p.i. (B) Histograms (top) and relative MFI (bottom) of indicated phenotypic markers in the SMARTA compartment 10 days p.i. (C) Identification of Th1 (Ly6C^hi^PSGL1^hi^) and Tfh (Ly6C^lo^PSGL1^lo^) subset in the SMARTA compartment by flow cytometry 10 days p.i. (D) Proportion of Tfh (left), Th1 cells (middle) and the Th1:Tfh ratio (right) of the SMARTA compartment 10 days p.i. (E) Histograms (left) and relative MFI (right) of Bcl6 and Tbet expression in the SMARTA compartment 10 days p.i (F) Bcl6 and Tbet MFI in SMARTA Th1 and Tfh subsets. Data are pooled from n = 2 independent experiments with 7-9 samples per group. Bars represent the mean and symbols represent individual mice. Significance was determined by unpaired two-tailed Student’s t-tests.

We next examined the phenotype of SMARTA T cells, focusing our analyses on effector cells due to the impaired generation of Tfh memory by SMARTA T cells(*26*). As the Y72F variant induced very few effector cells, we excluded this strain from further investigation. Consistent with earlier reports, strong T cell stimulation induced a larger proportion of Ly6c^+^ Th1 effectors, whereas the proportion of Tfh effectors was decreased (Figure 2B-D)(*4, 8-10*). In contrast, the ratio of Th1 and Tfh effector cells generated by host NP-specific T cells was consistent across all viral strains, providing an internal control for the comparable ability of these viruses to induce endogenous T cell responses (Figure 2D, Figure S2A). Expression of folate receptor 4 (FR4), an alternative marker for Tfh cell identification, was additionally used in combination with PD1 to discriminate the Tfh cell compartment, and demonstrated a decreased proportion of Tfh cells activated by strong compared to weak TCR stimulation (Fig. S2B)(*26, 27*). Accordingly, the expression of Bcl6 and T-bet, lineage defining transcription factors for Tfh and Th1 cells, respectively, revealed a mild but significant trend toward increased Tbet and decreased Bcl6 in response to strong stimulation (Fig. 2E). Importantly, Bcl6 expression was higher on Tfh compared to Th1 effectors, with no differences observed between strong and weak stimulation (Figure 2F, S2C). These data indicate that TCR signal strength is unlikely to exert a qualitative impact on these subsets. Notably, we did not observe any impact of TCR signal strength on the development of PSGL1^hi^Ly6c^lo^ T cells, previously reported to be a less differentiated population of Th1 effectors (Figure S2D)(*26*). In sum, consistent with earlier reports, TCR signal strength positively correlates with an increased ratio of Th1 to Tfh effectors during acute LCMV infection.

**Figure S2:**
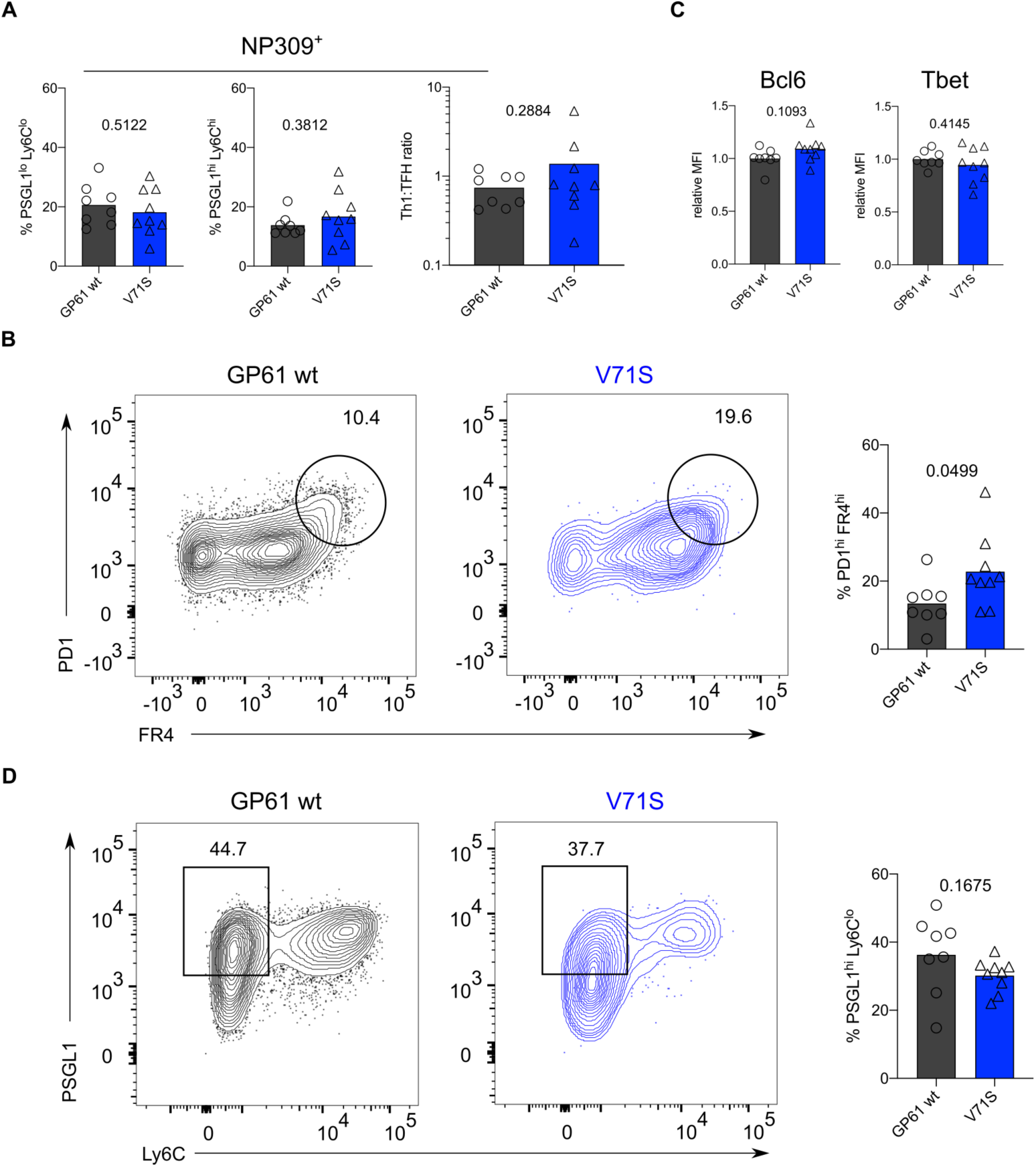
TCR signal strength positively correlates with Th1 cell differentiation during LCMV Armstrong variant infection. (A) Proportion of Tfh (left), Th1 cells (middle) and the Th1:Tfh ratio (right) of the NP309^+^ compartment 10 days p.i. (B) Identification and proportion of PD1^hi^ FR4^hi^ Tfh cells in the SMARTA compartment by flow cytometry. (C) Relative MFI of Bcl6 and Tbet expression in the Ly6C^lo^ Th1 SMARTA compartment. (D) Identification and proportion of Ly6C^lo^ Th1 (Ly6C^lo^PSGL1^hi^) in the SMARTA compartment by flow cytometry. Data are pooled from n = 2 independent experiments with 8-9 samples per group. Bars represent the mean and symbols represent individual mice. Significance was determined by unpaired two-tailed Student’s t-tests.

### TCR signal strength positively correlates with Tfh cell differentiation during LCMV Clone-13 variant infection

In contrast to acute LCMV infection, SMARTA T cells responding to chronic LCMV preferentially adopt a Tfh effector phenotype(*14, 15*). The impact of TCR signal strength within this context has not been determined, although affinity diversity among endogenous T cells is reportedly similar between acute and chronic LCMV infection(*28*). To directly assess the impact of TCR signal strength during chronic infection we transferred SMARTA T cells into congenic recipients followed by infection with rLCMV Clone-13 expressing either GP61 wt, V71S or Y72F. As an additional control we infected mice with rLCMV Armstrong which induced a similar expansion of SMARTA, NP-specific and CD4^+^CD44^+^ T cells as its Clone-13 counterpart (Fig S3A). Consistent with the results from acute infection, SMARTA T cell numbers at day 7 p.i. positively correlated with TCR signal strength, while the expansion of NP-specific and CD4^+^CD44^+^ T cells was similar in response to all three Clone-13 variants (Figure 3A). Importantly, infection with rLCMV Clone-13 Y72F induced approximately 2-fold more SMARTA T cell effectors compared to acute infection, allowing for a thorough investigation of T cells responding to this very weak potency variant (Figure 3A, Figure 2A).

**Figure 3:**
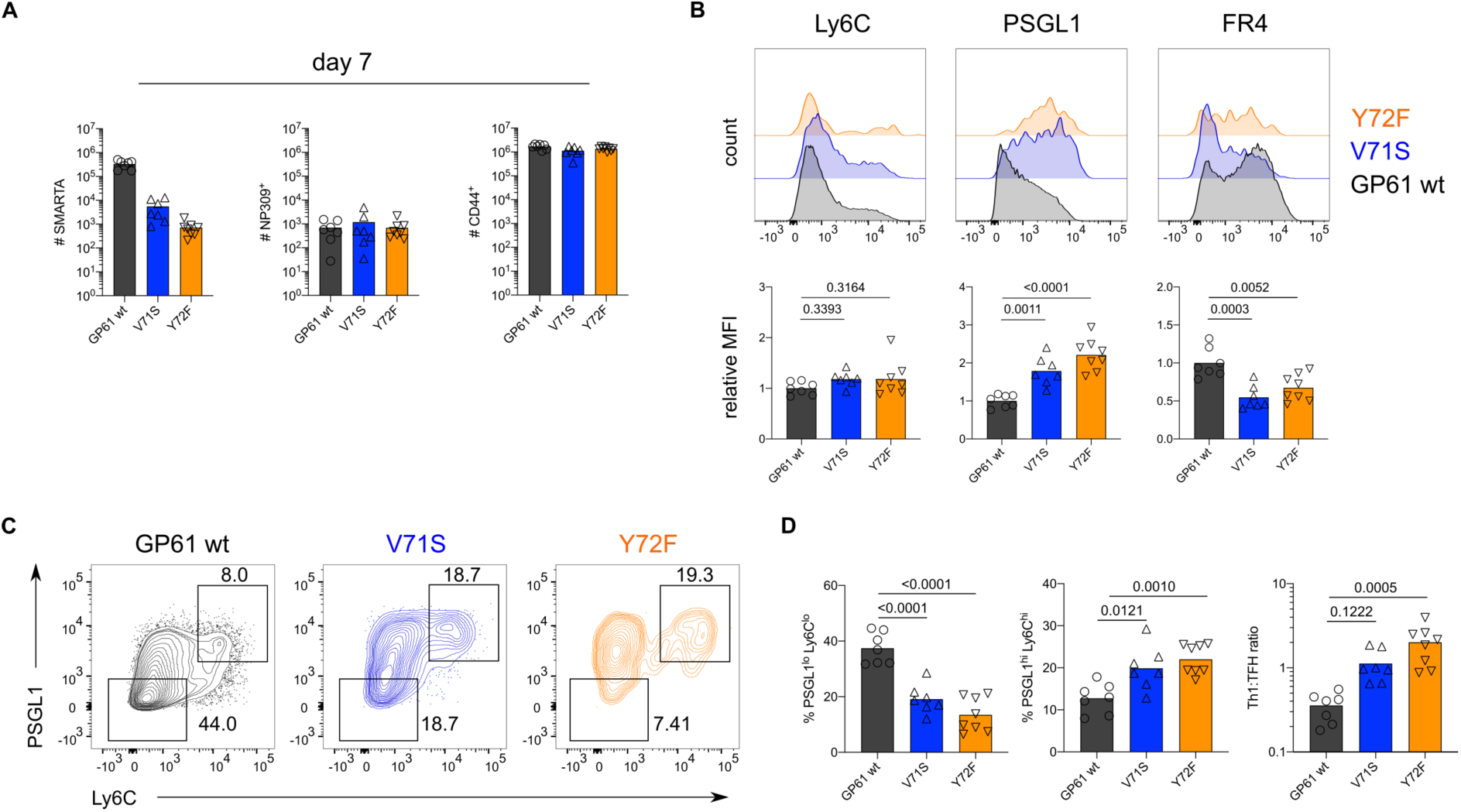
TCR signal strength positively correlates with Tfh cell differentiation during LCMV Clone-13 variant infection. Spleens were harvested 7 days after infection with LCMV Clone-13 variants. (A) Number of SMARTA (left), NP309^+^ (middle) and CD44^+^ cells (right). (B) Histograms (top) and relative MFI (bottom) of indicated phenotypic markers in the SMARTA compartment. (C) Identification of Th1 (Ly6C^hi^PSGL1^hi^) and Tfh (Ly6C^lo^PSGL1^lo^) subset in the SMARTA compartment by flow cytometry. (D) Proportion of Tfh (left), Th1 cells (middle) and the Th1:Tfh ratio (right) of the SMARTA compartment. Data are pooled from n = 2 independent experiments with 7-8 samples per group. Bars represent the mean and symbols represent individual mice. Significance was determined by one-way analysis of variance (ANOVA) followed by Tukey’s post-test.

With respect to T cell phenotype, strong TCR stimulation during rLCMV Clone-13 GP61 wt infection shifted the balance toward Tfh effector cell differentiation when compared to strong TCR stimulation in the context of acute infection (Figure S3B-C). Unexpectedly, and in contrast to the Armstrong variants, weaker TCR signaling during Clone-13 variant infection resulted in increased proportions of both PSGL1^hi^Ly6c^hi^ and PSGL1^hi^Ly6c^lo^ Th1 cells with the weakest variant, Y72F, generating the highest proportion of Th1 effectors (Figure 3B-D, S3D). The shift toward Th1 effectors in response to lower TCR signal strength is unlikely due to differences in antigen load as all variants sustained high viral titers in the kidneys at day 7 p.i (Figure S3E). In addition, although the viral titer of intermediate potency variant V71S was slightly decreased compared to GP61 wt and Y72F infection, NP-specific CD4 T cells exhibited a similar ratio of Th1 to Tfh effectors across all three infections (Figure S3F). Taken together, these reveal that TCR signal strength differentially modulates T cell fate acquisition according to the infectious context.

**Figure S3:**
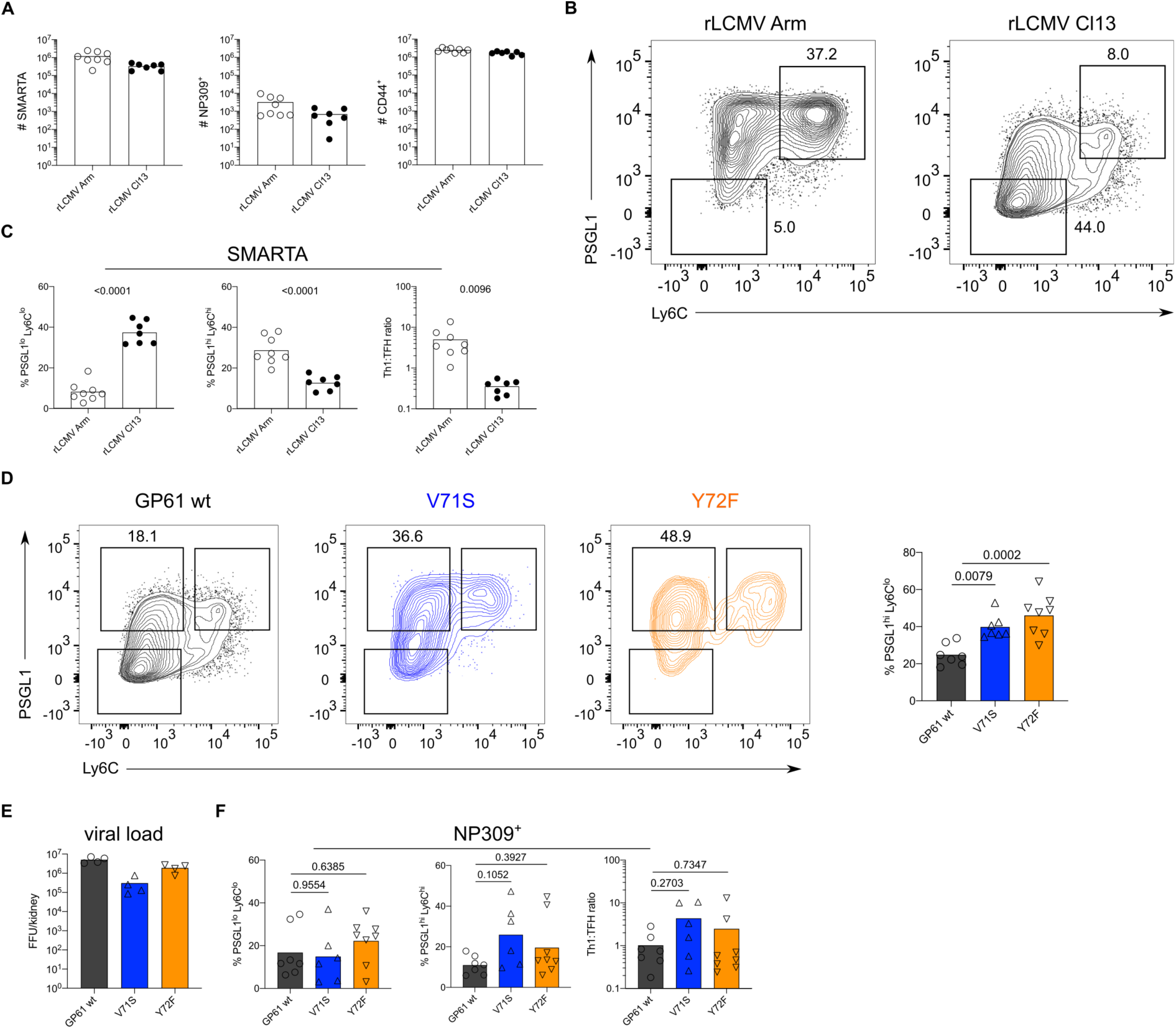
TCR signal strength positively correlates with Tfh cell differentiation during LCMV Clone-13 variant infection. Spleens were harvested 7 days after infection with LCMV Clone-13 variants. (A) Number of SMARTA (left), NP309^+^ (middle) and CD44^+^ cells (right). (B) Identification of Th1 (Ly6C^hi^PSGL1^hi^) and Tfh (Ly6C^lo^PSGL1^lo^) subset in the SMARTA compartment by flow cytometry. (C) Proportion of Tfh (left), Th1 cells (middle) and the Th1:Tfh ratio (right) of the SMARTA compartment. (D) Identification and proportion of Ly6C^lo^ Th1 (Ly6C^lo^PSGL1^hi^) in the SMARTA compartment by flow cytometry. (E) Viral load in kidneys. (F) Proportion of Tfh (left), Th1 cells (middle) and the Th1:Tfh ratio (right) of the NP309^+^ compartment. Data are pooled from n = 2 independent experiments with 7-8 samples per group except for (E) where one representative experiment of n = 2 independent experiments is shown with 4 samples per group. Bars represent the mean and symbols represent individual mice. Significance was determined by one-way analysis of variance (ANOVA) followed by Tukey’s post-test.

### Increasing TCR signal strength promotes the expression of markers associated with chronic T cell stimulation

During Clone-13 infection, T cells start to upregulate inhibitory surface markers associated with chronic activation, a state often referred to as “exhaustion”(*15, 29-31*). To understand if TCR signal strength impacts the expression of these markers we analyzed SMARTA T cells responding to Clone-13 GP61 wt and variant viruses at day 14. T cells responding to strong TCR signals expressed the highest levels of both PD1 and Lag3, two well characterized co-inhibitory receptors (Figure 4A-B) (*15, 29*). SMARTA T cells co-expressing both PD1 and Lag3 were most abundant following Clone-13 GP61 wt infection and decreased in response to Clone-13 variant infection (Figure 4C-D). Although the viral load was decreased in Clone-13 variant infections at this time point, the basal activation of CD4^+^CD44^+^ T cells was equivalent across all three strains and clearly above the recombinant LCMV Armstrong control (Figure S4A-B). Next, we examined the expression of TOX, a transcription factor involved in the adaptation of CD8 T cells to chronic infection(*32-36*). In response to acute infection, SMARTA Tfh cells expressed higher levels of TOX compared to Th1 cells, consistent with an earlier study highlighting the importance of TOX for Tfh cell development (Figure S4C)(*37*). In contrast, TOX expression during rLCMV Clone-13 wt infection was most highly upregulated by Th1 effectors (Figure S4C). In line with the expression of PD1 and Lag3, TOX was decreased on SMARTA T cells responding to rLCMV Clone-13 variant viruses, despite being comparably induced on CD4^+^CD44^+^ T cells (Figure 4E, Figure S4D). TOX was recently demonstrated to be important for the survival of stem-like TCF1^+^ CD8 T cells that accumulate during chronic LCMV(*34, 38, 39*). Given the transcriptional similarities of TCF1^+^ CD8 T cells and Tfh cells, we wondered if TCF1 would be similarly regulated by TCR signal strength following Clone-13 variant infection(*40*). Here we observed that unlike TOX expression, TCF1 is similarly expressed by T cells responding to all three rLCMV Clone-13 variants (Figure 4F, S4E), indicating that TCF1 expression is likely to be maintained independently of TCR signals.

**Figure 4:**
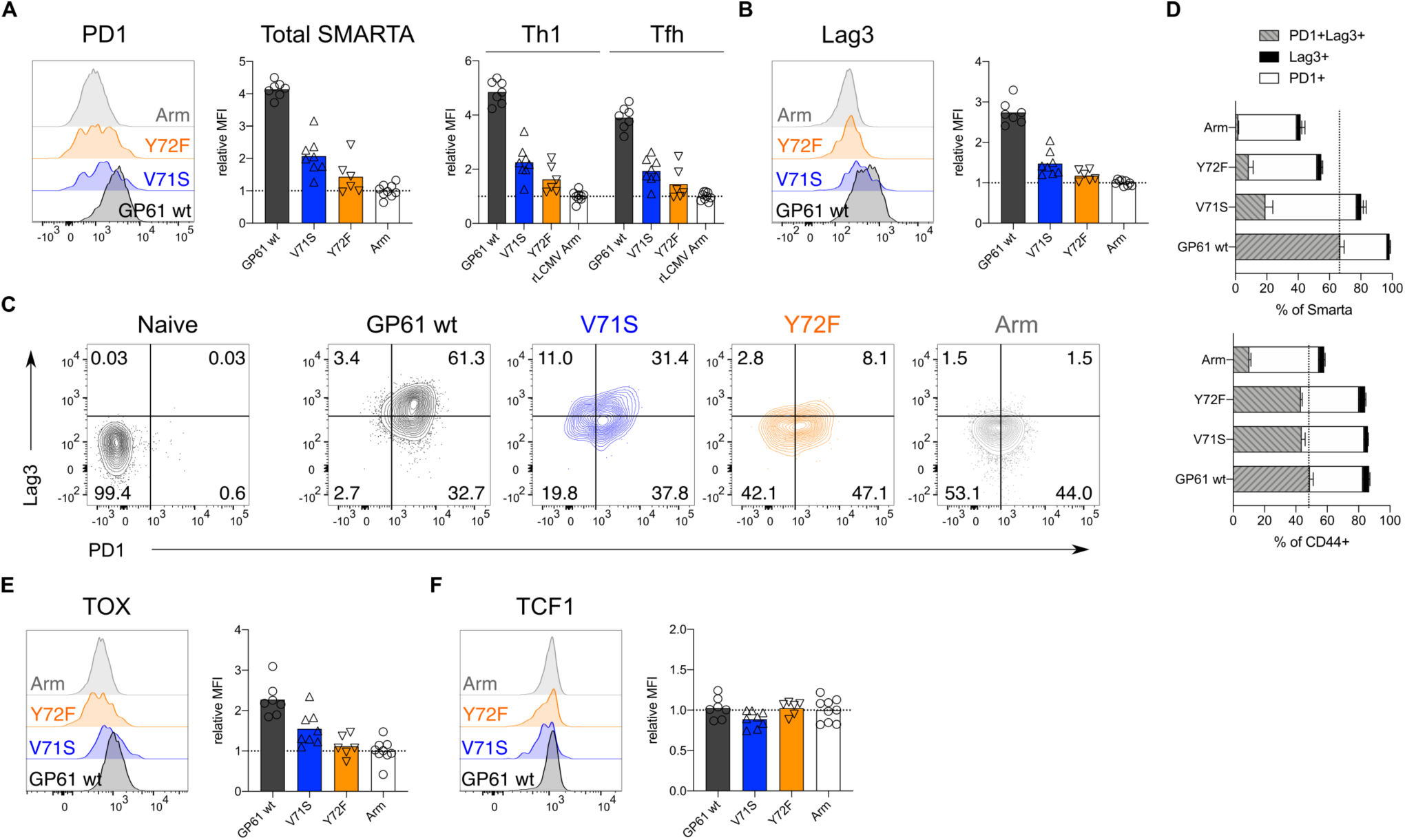
Increased TCR signal strength induces expression of markers associated with chronic T cell stimulation. Spleens were harvested 14 days after infection with LCMV Clone-13 based variants. (A) Histograms (left) and relative MFI (right) of PD1 in the total SMARTA compartment (left) or SMARTA Th1 and Tfh subsets (right). (B) Histograms (left) and relative MFI (right) of Lag3 in the SMARTA compartment. (C) Identification of PD1^+^Lag3^+^ SMARTA cells by flow cytometry compared to naïve CD62L^+^ CD44^-^ CD4 T cells from an uninfected mouse. (D) Quantification of PD1^+^Lag3^+^ SMARTA cells in the SMARTA (top) or CD44^+^ (bottom) compartment. (E) Histogram (left) and relative MFI (right) of TOX in the SMARTA compartment. (F) Histogram (left) and relative MFI (right) of TCF1 in the SMARTA compartment. Data are pooled from n = 2 independent experiments with 6-9 samples per group. Bars represent the mean and symbols represent individual mice. Significance was determined by one-way analysis of variance (ANOVA) followed by Tukey’s post-test.

**Figure S4:**
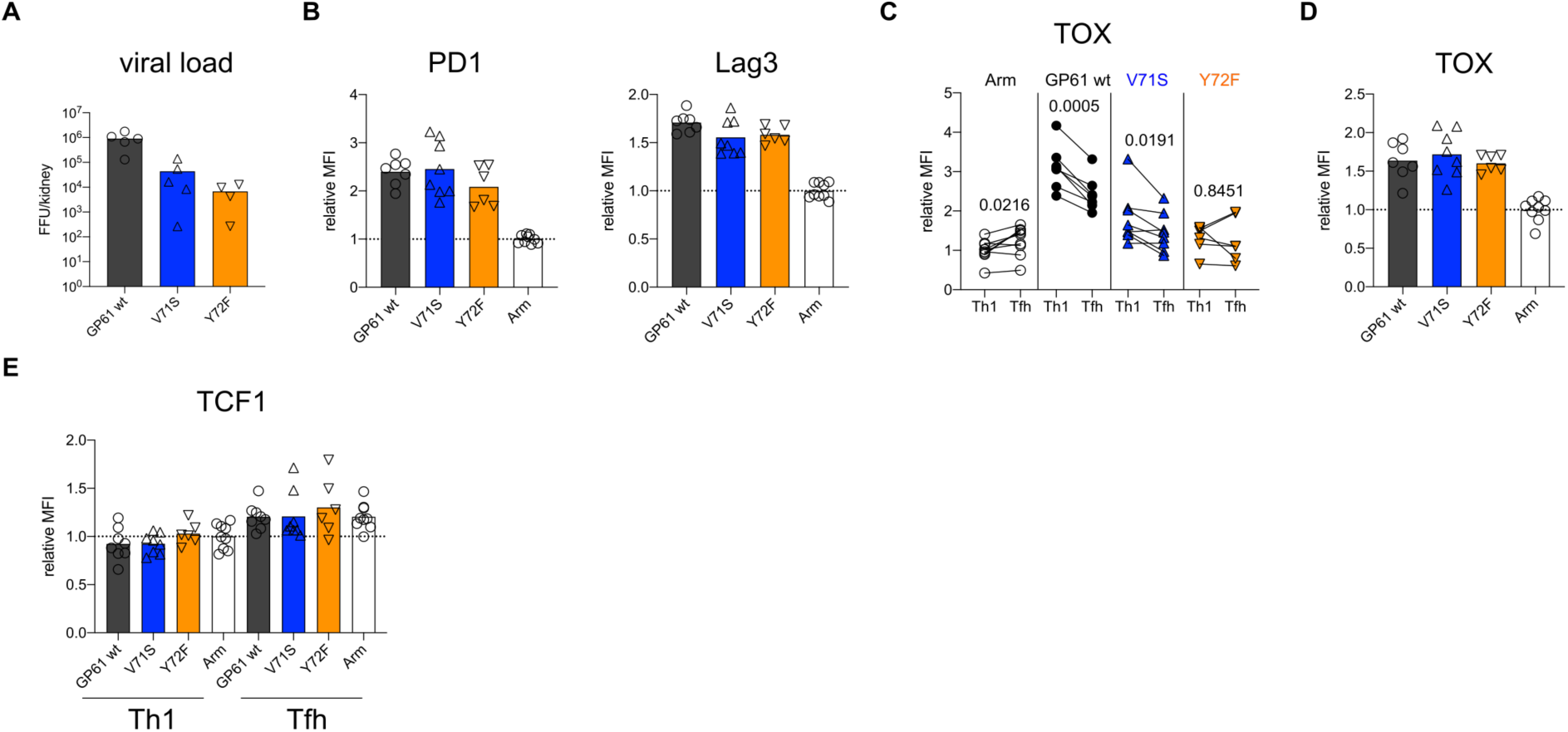
Increased TCR signal strength induces expression of markers associated with chronic T cell stimulation. Spleens were harvested 14 days after infection with LCMV Clone-13 based variants. (A) Viral load in kidneys. (B) Relative MFI (right) of PD1 and Lag3 in the CD44^+^ compartment. (C) Relative MFI of TOX in SMARTA Th1 and Tfh. (D) Relative MFI (right) of TOX in the CD44^+^ compartment. (E) Relative MFI of TCF1 in the SMARTA Th1 and Tfh subsets. Data are pooled from n = 2 independent experiments with 6-9 samples per group except for (A) where one representative experiment of n = 2 independent experiments is shown with 4-5 samples per group. Bars represent the mean and symbols represent individual mice. Significance was determined by one-way analysis of variance (ANOVA) followed by Tukey’s post-test (B,D) or by paired two-tailed Student’s t-tests (C).

## Discussion

The results of this study highlight the differential impact of TCR signal strength in shaping CD4 T cell fate according to the infection context. By systematically comparing the differentiation of TCR transgenic T cells responding to variant ligands in two distinct infection models, we demonstrate that the impact of TCR signal strength is heavily dependent on the infection specific parameters such as antigen load and inflammation.

The observation that TCR signal intensity correlates with Th1 generation during acute infection is consistent with accumulating evidence that higher potency ligands increase T cell sensitivity to IL-2, which likely drives the survival and expansion of Th1 effectors (*4, 8, 41-45*). This is similar to the paradigm described for Th1/Th2 cell differentiation, where stronger signals induce Th1 cells and weaker signals induce Th2 cells(*46*). Nevertheless, at very high antigen doses, T cells revert to Th2 differentiation, potentially due to the susceptibility of Th1 cells to activation induced cell death (AICD)(*47, 48*). AICD of Th1 cells might also contribute to biased Tfh generation at the higher end of TCR signal strength during Clone-13 infection(*49, 50*).

The shift of relatively high affinity CD4 T cells toward a Tfh cell phenotype during Clone-13 infection is well documented(*14, 40, 51*). In addition to antigen persistence, however, Clone-13 presents an altered inflammatory environment which contributes to an interferon stimulated gene signature and IL-10 production by chronically activated CD4 T cells(*15, 52*). It is possible that the unique cytokine milieu present during Clone-13 infection cooperates with strong TCR signals to fine tune T cell fate. For example, activation of T cells in the presence of IFNα induces T cell secretion of IL-10 which is positively regulated by TCR signal strength(*53, 54*). While this may ultimately serve to limit host pathology, it may also prevent the accumulation of Th1 effectors. Consistent with this idea, blocking IFNα or IL-10 during Clone-13 infection rescues the Th1 effector compartment and improves viral control, although this likely depends on the rate of viral replication(*55-57*). Within the same inflammatory context, weaker TCR signals might induce less T cell derived IL-10 which has been shown to impair Th1 effector cell differentiation(*52*). Of particular interest, T cell production of IL-10 during chronic infection depends on sustained, but ERK-independent TCR signals, suggesting that inflammatory versus suppressive cytokine secretion may have distinct TCR signaling requirements(*52*). Future experiments should address this by determining whether TCR signal strength contributes to cytokine production as well as cytokine susceptibility (i.e. induction of cytokine receptors) of effector cells responding during acute and chronic viral infection.

Finally, the ability of weakly activated T cells to maintain a higher proportion of Th1 effectors might ultimately contribute to viral control. The observation that the weakest Clone-13 variant, Y72F, elicited significantly more expansion than its Armstrong counterpart demonstrates that prolonged antigen presentation supports the accumulation of relatively low affinity T cells. Importantly, our study only follows the differentiation of T cells specific for a single epitope, while low affinity T cells are demonstrated to comprise up to half of the endogenous effector T cell response(*58*). Going forward, it will be interesting to determine whether targeting the expansion of lower affinity T cells with the potential to resist functional inactivation and maintain proliferative potential will improve control of viral infection.

## Acknowledgments

We thank all members of the Pinschewer lab for helpful discussion and David Schreiner for editing of the manuscript. Research was supported by the Swiss National Science Foundation (SNF, grants number PP00P3_157520 to CGK and number 310030_173132 to DDP), Gottfried and Julia Bangerter-Rhyner Stiftung, Olga Mayenfisch Stiftung, the Nikolaus and Bertha Burckhardt-Bürgin Stiftung (NBB), and the Freiwillige Akademische Gesellschaft (FAG) Basel.

## Author contributions

C.G.K. conceptualized the project. M.K. and C.G.K. designed the experiments, analyzed the data, wrote the manuscript and acquired funding. M.K. and P.R. performed experiments. D.P. acquired funding, provided advice on experimental design and revised the manuscript.

## Material & Methods

### Viruses

Virus rescue was performed as described previously using the pol-I/pol-II-driven reverse genetic system for LCMV (*21*). Single amino acids changes of the GP61-epitope were introduced by site directed mutagenesis of the previously described pI-S-WE-GP rescue plasmid (*21*). This plasmid encodes the nucleoprotein (NP) of the LCMV Armstrong strain *on cis* with the glycoprotein (GP) of LCMV WE. Additionally, the LCMV Armstrong specific D63K mutation was introduced into the GP61-coding sequence of the WE-GP gene matching the LCMV Armstrong / Clone-13 amino acid sequence of the GP61 peptides employed in the T cell activation assay. The resulting S-rescue plasmids were combined either a plasmid expressing either the Armstrong or the Clone-13 L segment in order to generate acute and chronic variants respectively. The presence of the desired mutations in the viral genomes was verified by sanger sequencing of RT-PCR amplicons generated with the OneStep RT-PCR-kit (Qiagen) using LCMV WE GP-specific primers (GATTGCGCTTTCCTCTAGATC and TCAGCGTCTTTTCCAGATAG). Viral RNA was extracted from cell culture supernatants using the Direct-zol RNA MicroPrep kit (Zymo Research). Virus titer were determined by immunofocus assay as described on NIH/3T3 cells (*59*).

### Viral growth kinetics

To determine viral replication capacities, BHK21 cells were seeded 24 hours prior to infection with a MOI of 0.01. Supernatant was collected at indicated time points and replaced with fresh culture medium.

### Mice and Animal experiments

Mice were bred and housed under specific pathogen-free conditions at the University Hospital of Basel according to the animal protection law in Switzerland. For all experiments, male or female sex-matched mice were used that were at least 6 weeks old at the time point of infection. The following mouse strains were used: C57BL/6 CD45.2, SMARTA Ly5.1, CD74^-/-^. Mice were injected with intraperitoneal injection of 2×10^5^ FFU for Armstrong variants or via intravenous injection of 2×10^6^ FFU for Clone-13 variants.

### NICD-protector

Mice were intravenously (i.v.) injected with 12.5μg homemade ARTC2.2-blocking nanobody s+16 (NICD-protector) at least 15 minutes prior to organ harvest.

### Adoptive cell transfer

Single-cell suspensions of cells were prepared from lymph nodes by mashing and filtering through a 100μm strainer. Naïve Smarta cells were enriched using Naïve CD4 T cell isolation kit (StemCell). 1×10^4^ SMARTA Ly5.1 cells were adoptively transferred into Ly5.2 recipients via intravenous injection as previously described (*60*).

### Flow Cytometry

Spleens were removed and single-cell suspensions were generated by mashing and filtering the spleens through a 100μm strainer followed by erythrocytes lysing using Ammonium-Chloride-Potassium (ACK) lysis buffer. SMARTA and endogenous LCMV-specific CD4 T cells were analyzed using IAb:NP309-328 (PE) or IAb:GP66-77 (APC) (provided by NIH tetramer core) tetramer. Following staining for 1 hour at room temperature in the presence of 50nM Dasatinib, tetramer-binding cells were enriched using magnetic beads and counted as previously published (*60*). Surface combined with viability staining was performed for 30min on ice. For transcription factor staining, fixation and permeabilization was performed according to the Foxp3/Transcription Factor staining kit (eBioscience). Samples were analyzed on Fortessa LSR II or Canto II cytometers (BD Biosciences) followed by data analysis with FlowJo X software (TreeStar). CD4^+^ T cells were pre-gated on lymphocytes in FSC/SSC, dump-, live CD4^+^ cells and then further gated on CD44^+^ Tetramer^-^ to assess the CD44^+^ compartment, CD44^+^ CD45.1^+^ GP66^+^ for SMARTA and CD44^+^ NP309^+^ for NP-specific cells.

The following antibodies were used: CD4 (BUV496, GK1.5, BD, #564667; APC eFluor780, GK1.5, eBioscience, #47-0041-82), CD11b (PE-Cy5, M1/70, BioLegend, #101222), B220 (PE-Cy5, RA3-6B2, BioLegend, #103210, PE-Cy7, RA3-6B2, BioLegend, #103222), CD11c (PE-Cy5, N418, BioLegend, #117316; AF647, N418, BioLegend, #117312), CD44 (BUV395, IM7, BD), CD45.1 (FITC, A20, BD, #553775; APC, A20, eBioscience, #17-0453-82), CD45.2 (APC-Fire, 104, BioLegend, #109852), CD69 (PE, H1.2F3, eBioscience), CD62L (APC, MEL-14, BD, #553152), PSGL-1 (BV605, 2PH1, BD, #740384), PD1 (BV785, 29F.1A12, BioLegend, #135225), Vα2 TCR (PE, B20.1, eBioscience, #12-5812-82; FITC, B20.1, eBioscience, #11-5812-82), Vβ8.3 TCR (FITC, 1B3.3, BD, #553663), F4/80 (PE-Cy5, BM8, BioLegend, #123112), I-Ab (PE, AF6-120.1, BD, #553552), FR4 (PE-Cy7, 12A5, BioLegend, #125012), Ly6C (BV510, HK1.4, BioLegend, #128033), Bcl6 (BV421, K112-91, BD, #563363), T-bet (BV711, 4B10, BioLegend, #644820), TOX (PE, TXRX10, eBioscience, #12-6502-82), Lag3 (BV421, C9B7W, BioLegend, #125221), TCF1 (AF700, 812145, R&D Systems, #FAB8224N), Zombie Fixable Viability Dye (Zombie Red, BioLegend, #423110).

### CD69 SMARTA activation assays

Serial dilutions of the GP61-wt peptide or APLs were plated. 5×10^5^ Ly5.2 Splenocytes and 1×10^5^ Ly5.1 SMARTA cells per well were added to the dilution series, stimulated overnight at 37°C, and subsequently stained and analyzed at the flow cytometer.

### MHC-II out-competition assays

CD74^-/-^ splenocytes were cultured with a custom made GP61-FITC at a fixed concentration of 1×10^−6^ M and various serial dilutions of GP61-wt or APLs for 4 hours at 37°C. After stimulation, the cells were stained and analyzed at the flow cytometer. The FITC-labelled GP61 peptide was custom made by Eurogentec.

### Statistical analysis

Geometric mean was used to determine the mean fluorescence intensity (MFI) and values were normalized to the mean of the control group from each experiment before data was pooled. Pooled and normalized MFIs are referred to as relative MFI. EC_50_ values were calculated using a sigmoidal dose-response fit in GraphPad Prism (version 8). For statistical analysis of one parameter between two groups, unpaired two-tailed Student’s t-tests were used to determine statistical significance. To compare one parameter between more than two groups, one-way analysis of variance (ANOVA) was used followed by Turkey’s post-test for multiple comparisons. P values are indicated on the graphs. Data was analyzed using GraphPad Prism software (version 8).

